# Adaptations accumulated under prolonged resource exhaustion are highly transient

**DOI:** 10.1101/2020.04.28.067322

**Authors:** Sarit Avrani, Sophia Katz, Ruth Hershberg

## Abstract

Many non-sproulating bacterial species can survive for years within exhausted growth media in a state termed long-term stationary phase (LTSP). We have been carrying out evolutionary experiments aimed at elucidating the dynamics of genetic adaptation under LTSP. We showed that *Escherichia coli* adapts to prolonged resource exhaustion through the highly convergent acquisition of mutations. In the most striking example of such convergent adaptation, we observed that across all independently evolving LTSP populations, over 90% of *E. coli* cells carry mutations to one of three specific sites of the RNA polymerase core enzyme (RNAPC). These LTSP adaptations reduce the ability of the cells carrying them to grow once fresh resources are again provided. Here, we examine how LTSP populations recover from costs associated with their adaptation, once resources are again provided to them. We demonstrate that due to the ability of LTSP populations to maintain high levels of standing genetic variation during adaptation, costly adaptations are very rapidly purged from the population once they are provided with fresh resources. We further demonstrate that recovery from costs acquired during adaptation under LTSP occurs much more rapidly than would be possible if LTSP adaptations had fixed during the time populations spent under resource exhaustion. Finally, we previously reported that under LTSP, some clones develop a mutator phenotype, greatly increasing their mutation accumulation rates. Here, we show that the mechanisms, by which populations recover from costs associated with fixed adaptations, may depend on mutator status.

## Introduction

Bacteria have a remarkable ability to rapidly adapt to almost any selective pressure applied on to them (Barrick and Lenski 2013). As they adapt, bacteria acquire costs associated with their adaptation (Cooper and Lenski 2000; Avrani et al. 2017). Such costs arise because mutations that are adaptive to one trait can often be harmful to other traits, a phenomenon termed antagonistic pleiotropy (Cooper and Lenski 2000; MacLean et al. 2004; Avrani et al. 2011; Kvitek and Sherlock 2013; Avrani et al. 2017). Additionally, as bacteria adapt to one condition they can accumulate mutations that are neutral under that condition but harmful once conditions shift (Cooper and Lenski 2000). The acquisition of costs associated with adaptation have been demonstrated under a variety of scenarios including but not limited to costs associated with the acquisition of resistance to antibiotics and phages (Andersson and Levin 1999; Reynolds 2000; Maisnier-Patin et al. 2002; Andersson and Hughes 2010; Conrad et al. 2010; Avrani et al. 2011; Avrani et al. 2017). Such costs have raised the question of whether adaptations will tend to persist once selection in their favor is no longer exerted.

Many studies have approached the question of whether bacterial populations fixed for adaptive mutations conferring resistance to an antibiotic or phage will tend to maintain resistance once no longer exposed to the antibiotic or phage (e.g. (Andersson and Levin 1999; Levin et al. 2000; Reynolds 2000; Maisnier-Patin et al. 2002; Andersson and Hughes 2010; Avrani and Lindell 2015)). Such studies have demonstrated that bacteria have a remarkable capability to compensate for the deleterious effects of resistance mutations by acquiring additional mutations. Since such compensatory mutations can occur at several different loci, while only one specific mutation can revert a resistance mutation back to its original non-resistant form, it was found that compensatory mutations tend to occur much more frequently than reversion mutations (Andersson and Levin 1999; Levin et al. 2000; Reynolds 2000; Maisnier-Patin et al. 2002; Andersson and Hughes 2010; Avrani and Lindell 2015). In other words, it was demonstrated that bacterial populations fixed for a resistance mutation much more frequently maintain their deleterious resistance allele by acquiring compensatory mutations, than lose their resistance through reversion mutations. Such studies seem to imply that costs associated with adaptation will rarely lead to reductions in the frequencies of adaptive alleles, once selection in their favor is no longer exerted.

Many non-sporulating bacterial species, including the model bacterium *Escherichia coli*, can survive for years within resource-exhausted media. When such bacterial species are inoculated into fresh media they undergo a short period of growth. Once their density increases beyond a certain level, they enter a short stationary phase followed by a period of rapid death in which viable cell numbers decrease by several orders of magnitude. However, some cells are able to survive death phase and enter a state termed long-term stationary phase (LTSP), in which populations can maintain fairly constant viable cell counts over many months and even years (Finkel and Kolter 1999; Finkel 2006; Avrani et al. 2017; Chib et al. 2017). By carrying out LTSP evolutionary experiments, followed up by whole genome sequencing of hundreds of evolved clones, we have previously characterized the dynamics of *E. coli* adaptation during the first four months (127 days) of survival under LTSP (Avrani et al. 2017). We have shown that *E. coli* can rapidly genetically adapt under LTSP in an extremely convergent manner, across independently evolving populations. In one of the most striking examples of such convergent adaptation, we found that 90% of bacteria surviving under LTSP carry a mutation within one of the three genes encoding the RNA polymerase core enzyme (RNAPC). In 93.5% of the clones carrying an RNAPC mutation, this mutation falls within one of only three specific sites: RpoB position 1272, rpoC position 334 or rpoC position 428. Clones carrying a mutation within each of these positions were found across all independently evolving sampled populations (Avrani et al. 2017). Such a striking pattern of convergent evolution clearly demonstrates that these RNAPC mutations are adaptive under LTSP. At the same time, we could also directly demonstrate that the RNAPC mutations are antagonistically pleiotropic as they reduce exponential growth rates within fresh rich media by ∼20% (Avrani et al. 2017). From now on we will refer to mutations falling within these three sites of the RNAPC as the antagonistically pleiotropic, convergent RNAPC adaptations, or anRNAPC for short.

In addition to the anRNAPC adaptations, LTSP populations acquire many additional mutations. In three of five sampled LTSP populations we observed the emergence of mutator clones, which acquired a mutation within a mismatch repair gene. Such clones carry higher numbers of mutations compared to non-mutator clones extracted at the same time point from the same or other populations. Mutations found within non-mutator clones seem to be mostly adaptive, while mutator clones tend to contain within them a higher fraction of apparently non-adaptive, passenger mutations (Avrani et al. 2017). Mutator clones carrying high numbers of mutations tended to suffer more severe reductions in their ability to grow within fresh LB. This indicates that, in addition to antagonistic pleiotropy, mutation accumulation during adaptation to LTSP also harms the ability of bacteria to grow once resources become available. Combined our results demonstrated that adaptation of *E. coli* under LTSP comes at a cost manifested in a reduced ability to grow once resources are again available.

Despite the fact that *E. coli* populations adapt rapidly in a highly convergent manner, suggesting very strong selection in favor of many of the observed adaptations, *E. coli* populations are able to maintain very high levels of genetic variation even as they adapt under LTSP.. While many adaptive mutations rise to very high frequencies, none of them fix across the entire population by day 127. The vast majority of clones sequenced from the day 127 LTSP populations carry relatively high numbers of mutations, including one of the convergent RNAPC mutations. However, we did observe some rare clones that had a genotype which was much more similar to their ancestral strain and did not carry a mutation within the RNAPC (Avrani et al. 2017).

Here we aimed to examine how LTSP adapted populations recover from costs acquired during the time they spent under resource exhaustion, once they are provided with fresh resources. We used LTSP adapted population samples and clones to compare and contrast the manner in which diverse adapted populations and individual adapted clones, fixed for specific genotypes, recover from costs associated with adaptation. We find that once LTSP adapted populations and individual clones are provided with fresh resources they can both rapidly recover their growth rates. However, population samples recover significantly more quickly than individual clones extracted from the same populations. Through whole genome sequencing of recovered clones we dissect the genetic mechanisms of such recovery. We can show that populations can recover through rapid fluctuations in allele frequencies, enabling them to select the least costly genotypes present within them from which to initiate recovery. This leads to rapid reductions in the frequencies of the most costly adaptations and to faster recovery than that observed for individual clones, fixed for specific genotypes. Furthermore, we show that even when recovery is initiated by a single clone, fixed for a specific costly adaptation, recovery will more often be achieved through reversion, if that clone is a mutator. Combined our results demonstrate that the dynamics of recovery from costs associated with adaptation are very different for populations in which diversity is maintained than for populations in which adaptive genotypes are fixed. Furthermore, our results indicate that due to their associated costs adaptive costly alleles may be far less persistent than largely thought.

## Results

### Whole population LTSP samples rapidly recover their growth rates within fresh media

Our LTSP evolutionary experiments were initiated by inoculating ∼5*10^6^ cells per ml of Luria Broth (LB) in a total volume of 400 ml within each of five 2 liter flasks. The five flasks were placed in an incubator shaker set to 37°C. Initially populations were sampled daily, then weekly, then monthly and finally at longer intervals. By plating dilutions of these samples, we could show that our populations entered LTSP at around day 11 of the experiment. ∼250 clones sampled at six time points spanning days 11 to 127 of the experiment were fully sequenced, alongside their ancestral clones, enabling us to reveal the dynamics of adaptation during this period of time (Avrani et al. 2017).

In the current study, we sought to examine how LTSP populations recover their ability to grow once provided with fresh resources. To do so, we carried out serial dilution experiments, starting with whole population samples extracted, from three of the five original LTSP populations (populations 1, 2, and 4. In order to maintain consistency with our previous publication, we will continue to refer thus to these populations here). In these experiments, three independent samples extracted from day 127 of each population were serially diluted 1:100 into fresh LB daily for 16 days. Given the 1:100 dilution, cells could grow for ∼6.6 generations per cycle. Similar serial dilution experiments were also carried out in parallel for four populations initiated with the ancestral *E. coli* K12 MG1655 (wildtype) strain used to initiate the original LTSP experiments. The different LTSP populations rapidly improved their ability to grow exponentially, relative the wildtype strain (Figure 1).

**Figure 1.**
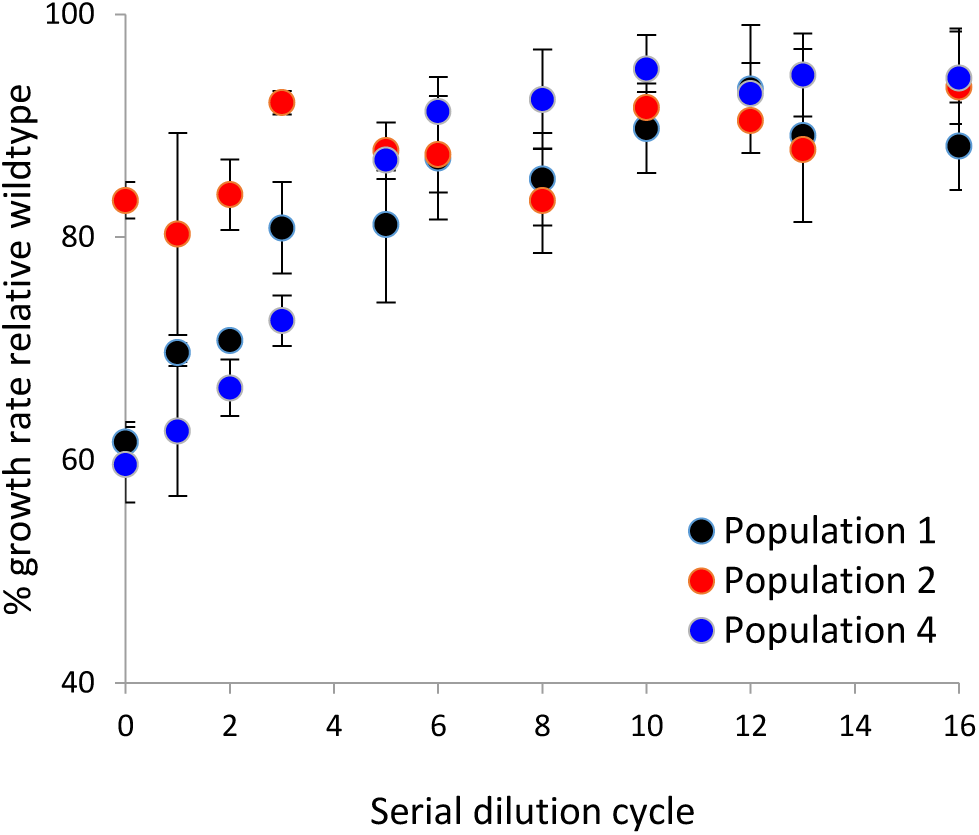
LTSP population samples rapidly recover their ancestral growth rates during serial dilution within fresh LB. The Y-axis represents the average growth rate relative the ancestral K12 MG1655 strain (wildtype), which was serially diluted alongside the population samples for a similar number of days. The average was calculated based on three independent serial dilution experiments for each of the three populations examined. The X-axis represents the number of serial dilution cycles within fresh LB that the population samples underwent. Error bars represent the standard deviation around the mean growth rate, relative the widltype.

### Recovery of whole population samples is associated with sharp reductions in the frequencies of antagonistically pleiotropic LTSP adaptive RNAPC alleles

In order to examine how the LTSP adapted populations recovered their growth rates, we fully sequenced ∼10 clones from one of the serial dilution experiments of each of the three populations following two, 10 and 16 days of serial dilution. A full list of the mutations identified is given in Table S1.

The original day 127 samples of all three populations all contained the anRNAPC adaptations at very high frequencies (ranging from 80% of sequenced clones in population 2 to 100% of sequenced clones in populations 1 and 4, Figure 2). By day 10 of serial dilution, across all populations we could no longer observe any anRNAPC adaptations (Figure 2). In population 1, these were largely replaced by other RNAPC mutations (Figure 2). These specific RNAPC mutations were observed within population 1 at earlier time points of the experiment (Avrani et al. 2017), indicating that they were likely also present, as standing variation in the day 127 samples of that population, at frequencies below our level of detection. In other words, it appears that in population 1 recovery was achieved through a fluctuation in genotype frequencies. This fluctuation led to the rapid replacement of the anRNAPC adaptation with another, initially less frequent, RNAPC adaptation.

**Figure 2.**
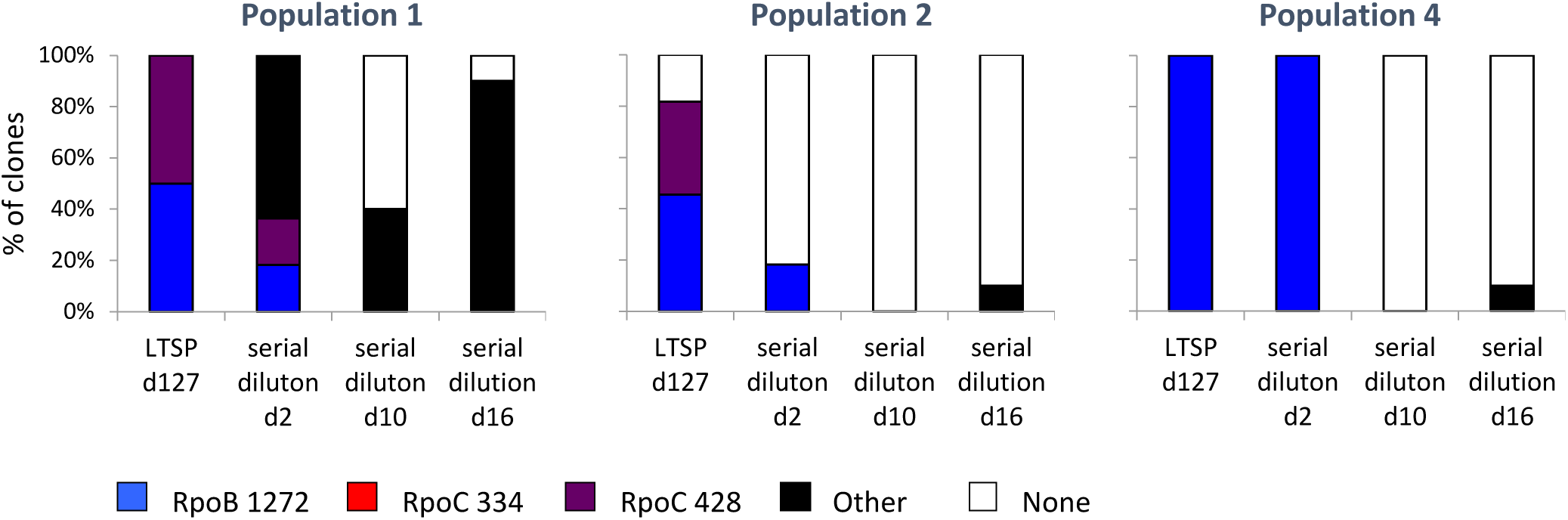
Recovery of ancestral growth rates of population samples is associated with the loss of the convergent antagonistically pleiotropic RNA polymerase core enzyme (anRNAPC) adaptations. Depicted are for each of the three populations the percentages of clones carrying each one of the three anRNAPC adaptations, another RNAPC mutation, or no RNAPC adaptation, immediately after sampling at day 127 of the original LTSP experiments and after two, 10 or 16 days of serial dilution.

Recovery in population 2 was also achieved from standing variation, through fluctuations in genotype frequencies. Population 2 was characterized by extremely high variation at day 127 of LTSP. While 80% of clones sequenced from population 2 at that time point carried an anRNAPC adaptation, 20% (two clones) did not (Avrani et al. 2017) (Figure 2). These two clones were in general very similar to the ancestral genotype as they carried a total of only 4 mutations each. The remaining sequenced clones varied greatly in their mutator status and in the number of mutations they carried. This variation manifested in substantial variation in each clone’s ability to grow within fresh LB (Figure 3). While some mutator clones with high mutation burden grew at only 30% of the ancestral exponential growth rates, the clones without anRNAPC adaptations were almost indistinguishable in their growth rate relative their ancestor. As a result, even at day 0 of serial dilution the entire population samples grew at ∼80% of the ancestral exponential growth rate (Figure 1, Figure 3). By day 2 of serial dilution the frequency of anRNAPC adaptations reduced to less than 20% (Figure 2). By day 10 clones carrying the anRNAPC adaptations were no longer detectable. Sequencing results further demonstrate that the clones that are observed during serial dilution that do not carry an anRNAPC adaptation are highly similar to the day 127 rarer clones that had only four mutations and did not have an anRNAPC adaptation (Table S1). Combined, these results clearly show that population 2 was able to rapidly recover from costs associated with adaptation under LTSP, through fluctuations in genotype frequencies. Such adaptation was possible due to the fact that population 2 had maintained extremely high levels of genetic variation during adaptation. This in turn allowed subsequent adaptation to regrowth to occur largely from standing variation, rather than through the occurrence of new compensatory or reversion mutations.

**Figure 3.**
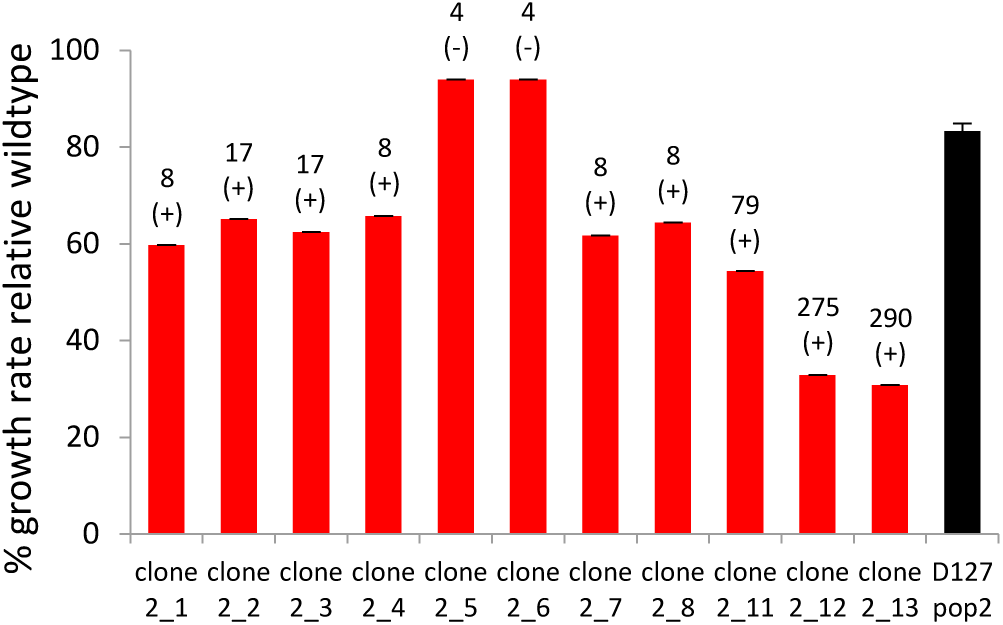
High variation in the exponential growth rates within fresh LB between clones extracted from LTSP population 2 at day 127 (day 0 of serial dilution). Depicted are the exponential growth rates of each clone relative that of the wildtype (red) and the mean exponential growth rate of four independent population 2 samples (black). The numbers above each red bar provide the number of mutations found within the corresponding clone. + signs indicate clones carrying an anRNAPC mutation and – signs indicate clones without an RNAPC mutation.

In population 4, recovery was also associated with sharp reductions in the frequencies of the anRNAPC adaptations. All clones sequenced from day 127 of LTSP population 4 were mutators carrying the same anRNAPC adaptation. The same is true for clones sequenced following two days of serial dilution. Clones sequenced from day 10 of serial dilution were also mutators, but no longer carried an anRNAPC adaptation. For this population it is hard to say whether the observed reductions in the frequencies of the anRNAPC adaptations resulted from fluctuations in genotype frequencies or from reversion mutations.

### LTSP adapted whole population samples recover their growth rates more rapidly than individual LTSP adapted clones extracted from the same samples

Next, we wanted to compare the ability of diverse LTSP adapted populations and individual clones, fixed for specific LTSP adapted genotypes to recover their ability to grow within fresh LB. Towards this end we focused on two of the three populations, population 2 and population 4. For these populations we initiated serial dilution experiments starting with each of the 10-11 clones sequenced from day 127 of those two populations (a total of 21 individual clones). Serial dilution experiments were carried out exactly as for the population samples above. As controls we also carried out, at the same time, serial dilutions using whole population samples from populations 2 and 4 and the ancestral wildtype strain. We found that while some clones were able to recover a close to ancestral growth rate, during the 16 days of the experiment, others were not able to do so (Table 1, Figure S1-S2). In population 2, all but three of the 11 clones examined recovered 90% or more of their ancestral growth rates (Table 1, Figure S1). In population 4 this was the case for only two of the 10 examined clones (Table 1, Figure S2). Even those clones that were successful in recovering their ancestral growth rate, almost always did so more slowly than the whole population samples of their population (Table 1, Figure 4, Figure S1 and Figure S2). The exceptions to this rule were the two clones observed in population 2 that carried only 4 mutations and did not carry any RNAPC mutations (Clones 2_5 and 2_6). These had to begin with suffered almost no cost to their ability to grow in fresh LB (Figure 3). It thus seems that the standing variation maintained within the LTSP populations enables them to recover their growth rates more rapidly, once provided with fresh resources, than is possible for most individual clones contained within them.

**Table 1.**
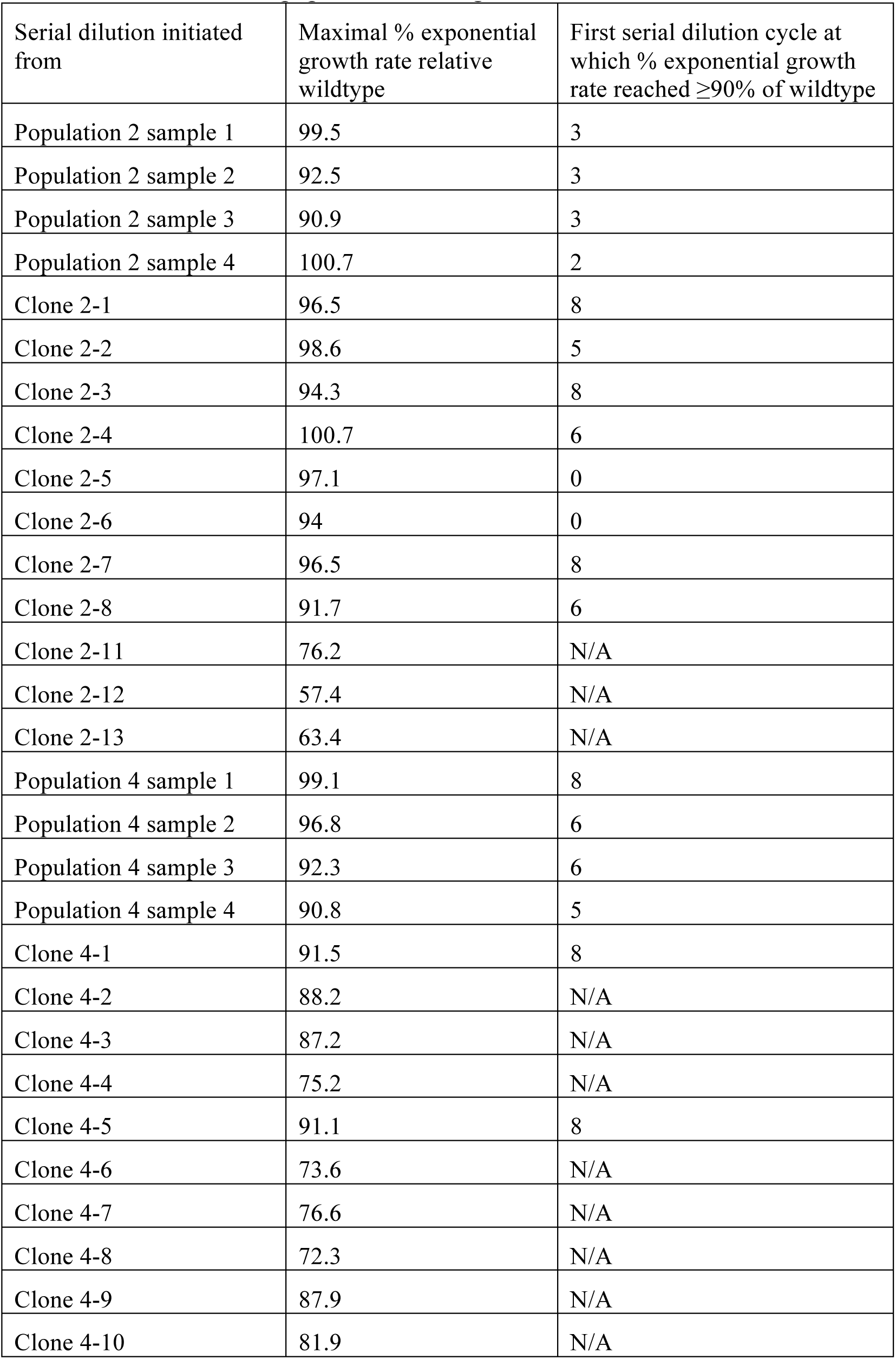
Recovery of LTSP day 127 population samples and individual clones extracted from the same populations during serial dilution into fresh LB

**Figure 4.**
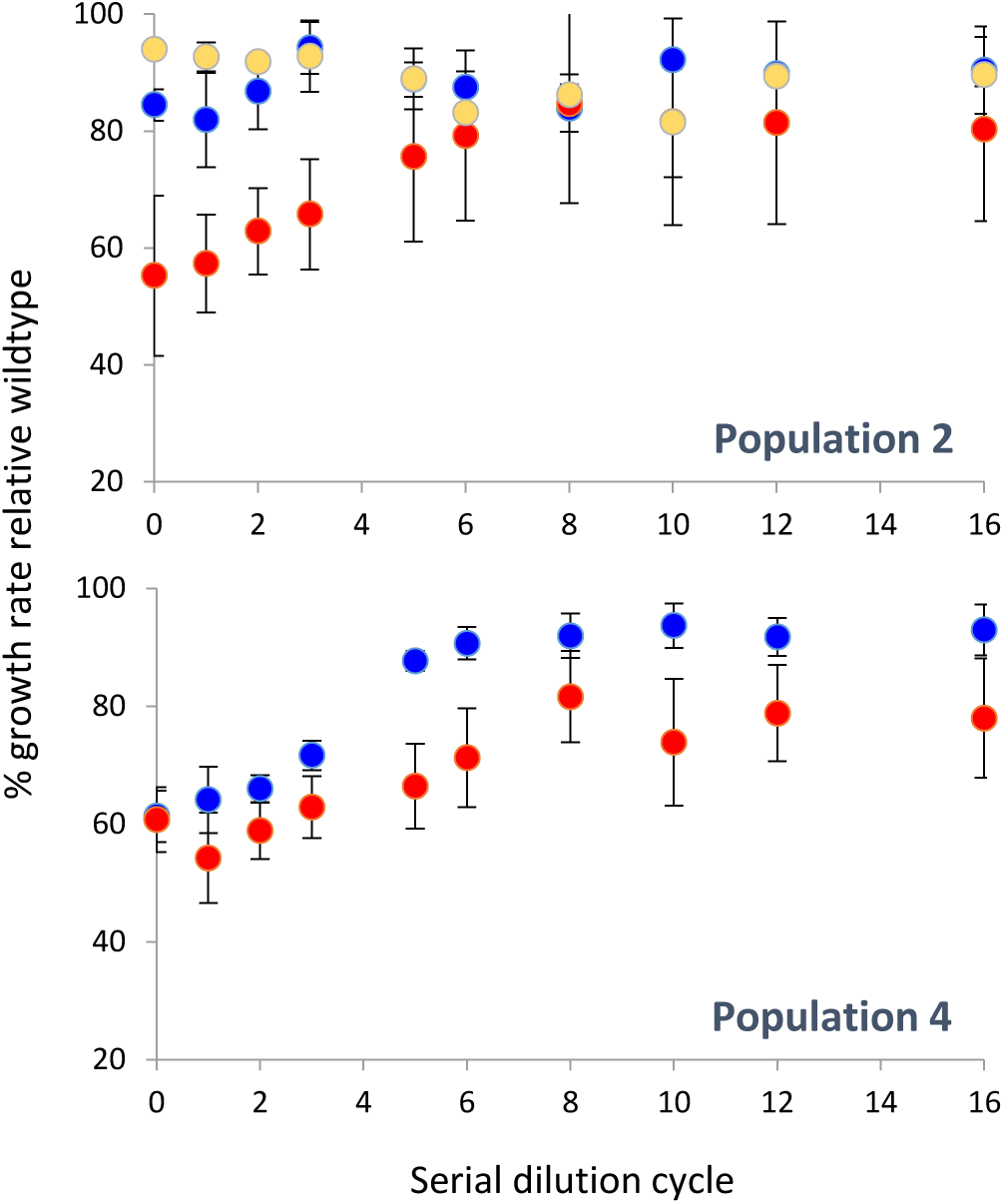
Whole population samples extracted following 127 days under LTSP, recover their growth rates within fresh LB, more rapidly than almost all individual LTSP clones extracted from the same time point and population. The results for populations 2 and 4 are presented separately. In both figures blue dots represent the mean growth rate of four independent population samples, relative the wildtype, which was serially diluted alongside the samples and clones, as a function of time (serial dilution cycles). The red dots represent the mean growth rate relative the wildtype, of all clones within that population that originally carried an anRNAPC mutation (nine of 11 clones for population 2 and 10 of 10 for population 4). In the population 2 figure, yellow dots represent the two clones that suffered only four mutations overall and did not suffer an RNAPC mutation. Error bars represent standard deviations around each mean.

### Mutator clones tend to recover from the costs of adaptation through reversion mutations

In order to examine how individual day 127 LTSP clones recover their ability to grow in fresh LB, we focused on four clones from population 2 and four clones from population 4. For each of these eight clones we sequenced ∼10 clones sampled from day 16 of their serial dilution experiments. The eight examined clones were selected to vary in their initial mutational load (Table 2). For consistency sake with our previous publication we maintain the designations given to each clone in that work. Two of the clones selected for analysis were non-mutators, one carrying a anRNAPC adaptation and one not. The remaining six clones were all mutators with varying numbers of mutations, ranging from nine to 290. While six of the examined clones managed to recover ≥90% of the ancestral wildtype exponential growth rate in fresh LB, two did not (Figure S1 and Figure S2). The two that did not (clones 2-13 and 4-4) were the ones with the highest mutation loads within their respective populations.

**Table 2.** Initial genotypes and numbers of acquired mutations for clones whose decedents were sequenced following 16 days of serial dilution

| Clone designation | Total # of mutations | Mutator status | anRNAPC mutation | Number of mutations acquired during 16 days of serial dilution |
| --- | --- | --- | --- | --- |
| 2_1 | 8 | Non mutator | Yes | 1-2 |
| 2_2 | 17 | Mutator | Yes | 27-45 |
| 2_6 | 4 | Non mutator | No | 2-4 |
| 2_13 | 290 | Mutator | Yes | 382-537 |
| 4_1 | 13 | Mutator | Yes | 27-39 |
| 4_4 | 24 | Mutator | Yes | 24-46 |
| 4_5 | 11 | Mutator | Yes | 17-32 |
| 4_9 | 9 | Mutator | Yes | 27-39 |

Since we are dealing here with individual clones that are initially fixed for a specific LTSP adapted genotype, recovery through fluctuation in initial genotype frequencies is not possible. Any observed loss of an anRNAPC allele must have therefore occurred via reversion mutations. From the sequencing results we could thus determine whether recovery was associated with the acquisition of compensatory mutations within the RNAPC, through reversion mutations or through a combination of both.

Clone 2-6 (4 mutations, no RNAPC mutation) did not really need to recover its growth rate as it was initially already very close to that of the wildtype ancestral genotype (Figure 3). Following 16 days of serial dilution clones arising from clone 2-6 acquired between two and four new mutations. These mutations were not enriched for non-synonymous mutations relative neutral expectations (Table 3), suggesting that most of them were likely not positively selected.

**Table 3.** Distribution of mutation types arising during serial dilution experiments

| Serial dilution initiated from | # total new mutations<sup>a</sup> | # non-synonymous new mutations<sup>a</sup> | # Synonymous new mutations<sup>a</sup> | dN/dS<sup>b</sup> | P-value<sup>c</sup> |
| --- | --- | --- | --- | --- | --- |
| Population 2 sample 1 | 41 | 14 | 5 | 0.86 | N/A |
| Population 2 sample 4 | 18 | 1 | 0 | N/A | N/A |
| Clone 2-1 | 12 | 10 | 0 | N/A | N/A |
| Clone 2-2 | 362 | 160 | 104 | 0.47 | <0.0001 |
| Clone 2-6 | 24 | 13 | 8 | 0.5 | N/A |
| Clone 2-13 | 4317 | 2259 | 1089 | 0.64 | <0.0001 |
| Population 4 sample 1 | 249 | 102 | 65 | 0.48 | <0.0001 |
| Population 4 sample 4 | 312 | 119 | 74 | 0.49 | <0.0001 |
| Clone 4-1 | 286 | 122 | 68 | 0.55 | <0.0001 |
| Clone 4-4 | 336 | 182 | 50 | 1.12 | 0.39 |
| Clone 4-5 | 252 | 106 | 86 | 0.38 | <0.0001 |
| Clone 4-9 | 322 | 160 | 70 | 0.7 | 0.017 |

<sup>a</sup>New mutations are mutations that did not appear in the ancestral clone with which the serial dilution experiments were initiated. In the case of whole population samples, new mutations are those mutations that were never seen in our sequencing of clones from that population at any of the six sequenced timepoints up to and including day 127. The sum of the non-synonymous and synonymous new mutations does not match the total number of new mutations, as mutations can also be non-coding, frame-shift, or nonsense.
<sup>b</sup>dN/dS (the ratio of the rates of non-synonymous to synonymous mutations) was calculated as
are given in the previous two columns of the table and number of synonymous and non-synonymous sites are calculated based on a combination of all *E. coli* protein-coding genes (see Materials and Methods)
<sup>c</sup> $\chi^2$ P-value with which it is possible to reject the null hypothesis that there is no enrichment in non-synonymous or synonymous mutations, relative random expectations based on the number of non-synonymous and synonymous sites within *E. coli* protein coding genes.

Clone 2-1 (non-mutator, anRNAPC mutation, eight mutations overall) managed to recover a close to ancestral growth rate by day eight of the serial dilution experiment (Figure S1, Table 1). From the sequencing results it is clear that this was achieved through the acquisition of a secondary compensatory mutation with the RNAPC. The 10 clones sequenced following 16 days of serial dilution all carried a secondary mutation within the RNAPC (Table S2). While each clone carried only one compensatory RNAPC mutation, we observed four different such mutations occurring within two sites of the RNAPC (RpoC position 1075 or RpoB position 546). Other than the compensatory RNAPC mutation, eight of the 10 sequenced clones suffered no additional mutations, while the remaining two clones suffered one additional mutation. Such a non-random pattern of mutation (mutations occurring specifically within two sites of the same gene as which the original antagonistically pleiotropic mutation was found and near to no additional mutations found) is a clear indication of compensatory adaptation.

The four mutator clones with the lower mutational burden that did manage to recover a near ancestral growth rate by day 16 of serial dilution, all did so by acquiring reversion mutations. This could be determined because the majority of clones sequenced from their day 16 populations no longer carried the anRNAPC mutation their ancestral clones originally carried (Table S2). In addition to these reversion mutations the clones suffered many additional mutations, likely due to them being mutators (Table 2). These mutations were not enriched for non-synonymous mutations relative random expectations, indicating that the majority of them were not adaptive mutations accumulated due to positive selection. If anything, the accumulated mutations were enriched for synonymous mutations (dN/dS significantly lower than 1, Table 3), indicating the removal of deleterious mutations from these populations by purifying selection. Populations evolved by serial dilution of two of these four clones (clone 4-1 and clone 4-9) acquired novel mutations within the RNAPC (Table S2). These mutations may be compensatory mutations that occurred prior to the occurrence of the reversion mutation that lead to the loss of the anRNAPC mutation. However further study will be needed to determine whether this is the case. The two mutator clones with the highest mutation loads within each of their original LTSP populations (clones 2-13 and 4-4) did not manage to recover a near-ancestral growth rate by day 16 of the serial dilution experiments (Figure S2). The sequencing results of these clones revealed that they acquired large numbers of mutations during serial dilution (Table 2). Mutations acquired by clone 4-4’s descendants were not significantly enriched for non-synonymous or synonymous mutations, relative random expectations (Table 3). Mutations carried by clones evolved from clone 2-13 were significantly enriched for synonymous mutations (Table 3), indicating evolution under purifying selection. In both the case of clone 4-4 and clone 2-13, all clones sequenced following 16 days of serial dilution, still carried their original anRNAPC mutations (Table S2).

Combined, our results showed that many, but not all clones extracted following 127 days of starvation are able to recover near ancestral growth rates following 16 days of serial dilution into fresh LB. However, even those clones that do recover well, tend to do so less rapidly than is possible for the entire population from which they were extracted. We further could show that while non-mutator clones may recover through compensatory evolution, leading to maintenance of the anRNAPC adaptations, mutator clones tended to recover through reversion mutations leading to loss of these antagonistically pleiotropic adaptations. Finally, mutator clones with high mutational loads could not fully recover their ancestral growth rates in 16 days of serial dilution. Unlike the descendants of lower-mutation-load mutator clones, the descendants of the high-load mutator clones did not lose the anRNAPC mutations they carried.

## Discussion

Our results demonstrate that due to the dynamics of adaptation under LTSP, LTSP adaptations are highly transient and rapidly reduce in frequency, once populations are again provided with fresh resources. These results may indicate that, more generally, rapid adaptations may tend to be far less persistent than previously thought. Several studies have shown that on the background of non-mutator genotypes, costly adaptations which are fixed across the entire population are less likely to revert to their ancestral genotype than to persist through the acquisition of compensatory adaptations (e.g. (Andersson and Levin 1999; Levin et al. 2000; Reynolds 2000; Maisnier-Patin et al. 2002; Andersson and Hughes 2010; Avrani and Lindell 2015)). In contrast to this trend, we show that fixed costly adaptations occurring within mutators tended to undergo reversion, once they were no longer favored. Mutators have been shown to be major players in driving rapid adaptation forward in numerous evolutionary experiments, and were shown to be abundant in more natural settings as well (e.g. (Matic et al. 1997; Sniegowski et al. 1997; Cooper and Lenski 2000; Denamur et al. 2002; Notley-McRobb et al. 2002; Labat et al. 2005; Pal et al. 2007; Avrani et al. 2017)). It is thus reasonable to predict that a substantial fraction of rapid adaptations occurs within such mutators and our results suggest that such adaptations might be far less likely to persist, once selection in their favor shifts. More studies will be needed to examine the general effect mutation rates may have on the tendency of fixed costly adaptations to persist, once selection in their favor shifts.

Most past studies aimed at understanding whether costly adaptations will persist, once they are no longer favored assumed that adaptations will always fix under strong selection. Recovery from costs associated with fixed adaptations can only be achieved through compensatory evolution or through reversion. Our results demonstrate that adaptations need not fix, even under strong and persistent selection. Instead, bacterial populations are able to maintain very high levels of standing genetic variation, including genotypes that do not contain the most costly adaptations. This enables the populations to recover from costs associated with adaptations through fluctuations in genotype frequencies, leading to very rapid reductions in the frequencies of costly adaptive alleles.

Evolutionary experiments have highlighted the remarkable capability bacteria have for rapid adaptation. An observation of many evolutionary experiments is that rapid adaptations often tend to occur through mutations to very central housekeeping genes (Hershberg 2017; Maddamsetti et al. 2017). The most obvious example of this trend is adaptations often occurring within one of the most central of all housekeeping genes, the RNA polymerase core enzyme (RNAPC). Mutations within RNAPC genes were shown to be involved in adaptation to a variety of selective pressures including high temperatures (Tenaillon et al. 2012), low nutrients (Conrad et al. 2010), exposure to ionizing radiation (Bruckbauer et al. 2019), and prolonged resource exhaustion (Avrani et al. 2017). Such prevalent rapid adaptation occurring within the most central housekeeping genes stands in apparent contrast to their particularly high levels of sequence conservation. Housekeeping genes in general and the RNAPC in particular tend to be extremely well conserved in their sequences. Indeed, conservation of the RNAPC is high enough to allow its genes to be used as slowly evolving markers for the study of bacterial phylogeny (Lan et al. 2016). How is it possible that RNAPC genes and other housekeeping genes would undergo very rapid sequence evolution at the short term in response to countless selective pressures, and yet stay so similar over longer evolutionary timescales? Our results may offer an explanation for this apparent discrepancy. Adaptations within housekeeping genes such as the RNAPC may occur quite frequently, but be highly transient due to their associated costs, leading to apparent high conservation over longer evolutionary distances.

Similar to other rapid adaptations, adaptations leading to antibiotic resistance also often occur within central housekeeping genes (including the RNAPC) and often come at a cost to fitness. In lab experiments involving lethal doses of antibiotics, resistance tends to always be fixed. After all, bacteria that are not resistant cannot survive exposure to lethal doses of antibiotics, within a flask or a tube. For this reason, in lab experiments examining how bacteria recover from costs associated with antibiotic resistance, researchers have most often focused on scenarios of fixed antibiotic resistance. This has lead researchers to conclude that costs associated with resistance will not tend to lead to sharp reductions in resistance frequencies, once treatment stops. However, within the bodies of hosts, resistance may increase in frequencies in response to antibiotic exposure, without becoming fixed, due to such phenomena as bacterial persistence and heterogeneous and incomplete antibiotic penetration (Whelton and Walker 1974; Balaban et al. 2004; MacLean and Vogwill 2015). Our results indicate that if resistance is not fixed, the dynamics of resistance maintenance may very well be highly affected by costs associated with antibiotic resistance adaptation. Furthermore, studies have shown that mutators may often contribute to the emergence of clinical antibiotic resistance (Macia et al. 2005; Mehta et al. 2019). Our results demonstrate that such mutators may be much more likely to lose, even fixed costly adaptations through reversion mutations, rather than maintain them through compensation. Further studies will be required in order to determine whether the patterns we observe recapitulate themselves when it comes to antibiotic resistance adaptations.

The fact that we observed that mutators tend to lose their costly adaptations through reversion mutations may indicate that compensatory mutations often do not fully compensate for all deleterious effects of a costly adaptation. After all, the difference between mutators and non-mutataors is that the mutators may acquire reversion mutations more rapidly due to higher mutational input. However, it is likely that in mutators, as in non-mutators, compensatory mutations will tend to occur prior to the occurance of reversion mutations (as there are several possible compensatory mutations and only a single possible reversion mutations). If there is no difference in the fitness of a clone carrying a costly adaptation together with a compensatory mutation alleviating its cost, and a clone that no longer carries the costly adaptation, there is no reason for mutator clones to undergo reversion, rather than compensation. Fitting with this, for two of the mutator clones we examined it appears that additional mutations occurred within the RNAPC prior to loss of their anRNAPC mutations. These may well be compensatory mutations that occurred prior to the reversion of the anRNAPC adaptations, but which did not fully alleviate the costs associated with these adaptations, which were later lost through reversion. More studies will need to be taken in order to determine the extent to which compensatory mutations alleviate costs associated with adaptation and whether such compensation is often incomplete, leading to eventual reversion of adaptations, even once some of their costs are alleviated.

Combined the results of this study demonstrate that due to the dynamics of adaptation under prolonged resource exhaustion, LTSP adaptations can be highly transient, rapidly reducing in frequency once populations are provided with fresh resources. In more general terms these results suggest that costs associated with adaptive mutations may often lead to sharp reductions in their frequencies, once they are no longer favored by selection.

## Materials and methods

### Recovery from LTSP

Three to four independent samples extracted from day 127 of each of the three populations (1, 2, and 4), as well as 21 individual clones isolated previously and fully sequenced (Avrani et al. 2017) from populations 2 and 4, were each inoculated into 4 ml of fresh LB, within 14 ml tubes and allowed to grow for an hour while shaking at 225 rpm in 37°C (these conditions were used throughout the study). These cultures were then transferred into 50 ml Erlenmeyer flasks containing additional 6ml of LB (for a total volume of 10 ml). These samples were each serially diluted 1:100 daily for 16 days. Similar serial dilution experiments were also carried out in parallel for populations initiated with the ancestral *E. coli* K12 MG1655 (wildtype) strain used to initiate the original LTSP experiments. Almost daily, a subsample of each culture was mixed with glycerol in a final concentration of 50% and then frozen in -80°C.

### Growth rate measurement

Prior to initiation of the experiments (day 0) and then almost daily during the experiments, a 100 µl sample of the diluted cultures were grown in a 96 well plate, in a plate reader. This plate was incubated in 37°C while shaking at 200 rpm and OD was measured every 10 minutes. The exponential growth rate was calculated from the slope of the growth at the exponential phase. We used the maximal rate calculated from five sequential time points, where R^2^ > 0.99. The calculated exponential growth rate was then compared to that of the ancestral *E. coli* K12 MG1655 (wildtype) strain, serially diluted along the LTSP population samples and clones to calculate the % growth rate relative the wildtype.

### Sequencing of the evolved clones

Frozen cultures of the desired populations and time points were thawed and dilutions were plated and grown over night. Ten colonies from each culture were used to inoculate 4 ml of medium in a test tube and were grown until they reached an OD of ∼1. This was done to reduce the time these colonies can evolve and accumulate mutations. 1 ml of the culture was centrifuged at 10,000 g for 5 min and the pellet was used for DNA extraction. The remainder of each culture was then archived by freezing in 50% of glycerol in a −80°C freezer. DNA was extracted using the MACHEREY-NAGEL Nucleo-Spin 96 Tissue kit. Library preparation followed the protocol outlined in (Baym et al. 2015). Sequencing was carried out at the Technion Genome Center using an Illumina HiSeq 2500 machine. All clones were sequenced using paired end 150 bp reads. The raw sequencing data will be deposited in full in the sequence read archive upon acceptance of the manuscript.

### Calling of Mutations

The reads obtained for each sequenced clone were aligned to the *E. coli* K12 MG1655 reference genome (accession NC_000913). The Breseq platform (Deatherage and Barrick 2014) was used for alignment and mutation calling. Breseq allows for the identification of point mutations, short insertions and deletions, larger deletions, and the creation of new junctions(Deatherage and Barrick 2014). All called mutations are provided in Tables S1 and S2.

### Calculating numbers of synonymous and non-synonymous sites within the *E. coli* K12 MG1655 genome

DNA sequences of all protein-coding genes of *E. coli* K12 MG1655 were downloaded from the NCBI database. The contribution of each protein-coding site to the count of non-synonymous and synonymous sites was calculated according to the likelihood that mutations to that site would lead to a non-synonymous or a synonymous changes. For example, mutations to the third codon position of a 4-fold degenerate codon would be 100% likely to be synonymous. Such a position would therefore add a count of 1 to the number of synonymous sites. In contrast mutations to the third codon position of a 2-fold-degenerate codon will be synonymous for a third of possible mutations and non-synonymous for two thirds of possible mutations. Such positions would therefore add a count of 1/3 to the number of synonymous sites and 2/3 to the number of non-synonymous sites. In such a manner, we could calculate what proportion of sites, across all *E. coli* K12 MG1655 protein-coding genes are non-synonymous (meaning that mutations to those sites would lead to non-synonymous changes) and what proportion are synonymous.

## Acknowledgments

This work was supported by an ISF grant (No. 756/17, to RH) and by the Rappaport Family Institute for Research in the Medical Sciences (to RH). The described work was carried out in the Rachel & Menachem Mendelovitch Evolutionary Process of Mutation & Natural Selection Research Laboratory.

## Figure legends

**Figure S1**. Whole population 2 samples extracted following 127 days under LTSP, recover their growth rates within fresh LB, more rapidly than almost all individual LTSP clones extracted from the same time point and population. Presented are individual graphs for the recovery of each of the clones, relative the recovery of the whole populations sample. The Y-axis represents the average growth rate relative the ancestral K12 MG1655 strain (wildtype), which was serially diluted alongside the population samples and individual clones for a similar number of days. The X-axis represents the number of serial dilution cycles within fresh LB that the population samples and individual clones underwent.

**Figure S2**. Whole population 4 samples extracted following 127 days under LTSP, recover their growth rates within fresh LB, more rapidly than almost all individual LTSP clones extracted from the same time point and population. Presented are individual graphs for the recovery of each of the clones, relative the recovery of the whole populations sample. The Y-axis represents the average growth rate relative the ancestral K12 MG1655 strain (wildtype), which was serially diluted alongside the population samples and individual clones for a similar number of days. The X-axis represents the number of serial dilution cycles within fresh LB that the population samples and individual clones underwent.

